# NLGN3 contributes to angiogenesis in myocardial infarction via activation of the Gαi1/3-Akt Pathway

**DOI:** 10.1101/2024.10.24.620138

**Authors:** Shunsong Qiao, Chao Tang, Jingjing Zhu, Cong Cao, Yu Feng, Xiaosong Gu

## Abstract

**BACKGROUND:** Angiogenesis is an important repair mechanism for myocardial infarction. Neuroligin-3 (NLGN3) can promote angiogenesis by activating Gαi1/3-Akt signaling following ischemic brain injury. This study investigated the role of NLGN3 in myocardial infarction (MI).

**METHODS AND RESULTS:** On the 7^th^ day after MI, the plasma level of NLGN3 in patients was significantly higher than in the control group. A mouse model of MI also showed significantly increased expression of NLGN3 in heart tissue. Single-nucleus transcriptome analysis revealed that NLGN3 was located predominantly in cardiac fibroblasts and endothelial cells (ECs). Endothelial-specific knockdown of NLGN3, or inhibition of NLGN3 using ADAM10i, significantly increased the ischemic area, reduced angiogenesis, and worsened cardiac function. Co-immunoprecipitation (Co-IP) experiments showed that NLGN3 interacted with Gαi1/3. The Gαi1/3 knockout (Gαi1/3-KO) mouse model of MI showed an increased ischemic area, decreased angiogenesis, and impaired cardiac function. Mechanistic studies showed that the NLGN3-Gαi1/3 signaling pathway exerts cardioprotective effects by promoting EC proliferation and tube formation through the PI3K-Akt-mTOR pathway. Silencing of Gαi1/3 largely eliminated the ability of NLGN3-promoting cardiac ECs to proliferate and form tubes.

**CONCLUSION:** Our findings suggest the endothelial NLGN3-Gαi1/3 signaling pathway promotes angiogenesis and reduces the ischemic area following MI, which is critical for maintaining cardiac function and repairing tissues. Targeting of the NLGN3-Gαi1/3 signaling pathway may have clinical therapeutic potential in protecting the heart from ischemic injury.

**What Is Known?:**

1. Myocardial infarction (MI) leads to a significantly increased risk of recurrent events in individually vascularized regions, which then stimulates adverse systemic vascular effects.
2. Gαi proteins play a key role in receptor tyrosine kinase (RTK) and other non-GPCR receptor-mediated signaling.
3. Neuroligin 3 (NLGN3) is a major member of the NLGN family of proteins that help to form synapses between neurons. It is known be cleaved and secreted in an activity-dependent manner, but its function in heart disease is still poorly understood.

**What New Information Does This Article Contribute?:**

1. NLGN3 expression was found to be markedly increased in the heart after MI, especially in cardiac ECs.
2. Knockdown of Gαi1/3 significantly reduced angiogenesis and impaired cardiac function, while knockdown of NLGN3 increased the ischemic area and reduced angiogenesis through Gαi1/3.
3. NLGN3-Gαi1/3 signaling promotes vascular EC proliferation and tube formation through the PI3K-Akt-mTOR pathway, thereby protecting the ischemic heart.

## INTRODUCTION

Myocardial infarction (MI) remains one of the principal causes of death and disability globally, and its treatment continues to pose a major challenge in contemporary cardiovascular medicine[1]. Approximately 10% of patients with MI survive with severely reduced left ventricular (LV) function, and such individuals are prone to develop myocardial remodeling[2]. Progressive myocardial remodeling occurs during the proliferative and reparative phase in which inflammation subsides, angiogenesis is induced, fibroblasts are activated, and fibrotic and scar tissue formation occurs[7]. Wound healing following MI involves a powerful angiogenic response that begins in the border zone and extends to the necrotic infarct core[8]. The promotion of angiogenesis in the infarcted area allows reperfusion and the preservation of surviving ischemic myocardium[9].

Angiogenesis is the morphogenesis of EC groaflumen surrounding a lesion through the association of individual ECs, or the sprouting of ECs from pre-existing vessels[5]. In traditional models of angiogenesis, ECs in the angiogenic ecotope respond to angiogenic signals by migrating toward the signal. The proximal majority of ECs then become tip cells (a transient EC state) and proliferate to form new vascular structures that differentiate into capillary (CapiECs), arterial (AEC), and venous (VEC) endothelial subtypes[6]. ECs are the largest population of non-cardiac myocytes in myocardial tissue[3]. Following MI, the process of myocardial repair is promoted by angiogenesis, which is dependent on the molecular biological regulation of ECs. Therefore, a better understanding of how ECs are regulated is valuable for protecting against the effects of MI.

Neural synaptic proteins are postsynaptic cell adhesion molecules expressed as four major isoforms: neuroligin-1 to -4, abbreviated as NLGN1 to NLGN4. These act as ligands for presynaptic neuroligins [10]. The synaptic protein neuroligin-3 (NLGN3) belongs to the family of neuroligins, a class of postsynaptic cell adhesion molecules that regulate synaptic organization and dendritic growth[11, 12]. NLGN3 can interact with neuroligins to mediate synaptic development and function, and has been linked to mental retardation syndromes[13]. Neuronal activity promotes the proliferation and growth of high-grade glioma by secreting NLGN3[14]. Researchers have identified a circRNA produced by the neuroligin (circNlgn) gene and which is upregulated in various congenital heart diseases with cardiac overload[15]. In addition, NLGN1 is known to cooperate with α6 integrins to direct EC interaction with the underlying extracellular matrix during vascular development, thus promoting angiogenesis[16]. NLGN3 is mainly cleaved by ADAM10 (A Disintegrin and Metalloproteinase 10) in neurons. Inhibition of ADAM10 was shown to prevent the cleavage of NLGN3 and its secretion into the microenvironment[17]. In a previous study, our group found that NLGN3 was significantly upregulated in mice with cerebral ischemia-reperfusion injury, where it exerted important protective effects[18]. However, the pathophysiologic function of NLGN3 in the heart is unknown. In the present study, we investigated whether NLGN3 was involved in remodeling after myocardial ischemic injury.

The three subunits of the inhibitory α-subunit of G protein (heterotrimeric guanine nucleotide binding protein), or Gαi protein, are Gαi1, Gαi2 and Gαi3[19]. By binding to GPCR (G protein-coupled receptor), Gαi proteins and βγ complexes inhibit adenylate cyclase (AC) and deplete cyclic AMP (cAMP) levels[19]. In addition, we showed that Gαi1 and Gαi3 proteins play key roles in mediating the signaling of several RTKs, including epidermal growth factor receptor (EGFR)[20], fibroblast growth factor receptor(FGFR)[21], keratinocyte growth factor receptor[22], and vascular endothelial growth factor receptor 2 (VEGFR2)[23]. Specifically, the recruitment of Gαi1 or Gαi3 subunits to RTKs following activation by ligands is essential for mediating PI3K-Akt-mTORC signaling[20-23]. Altered levels of Gαi protein expression have been found in heart disease, with human heart failure associated with the upregulation of Gαi2 and Gαi3[24]. By overexpressing constitutively active inhibitory Gαi proteins, gene therapy-based modulation of atrioventricular conduction effectively reduced the heart rate in ventricular fibrillation (AF)[25]. Furthermore, Gαi signaling reduces myocardial infarct size, inhibits ischemia-induced apoptosis, and in some cases enhances the recovery of contractile function after ischemia[26, 27]. We previously reported that NLGN3 can regulate the growth of nerve cells through the Gαi1/3 pathway[43]. In the present study we explored the potential function of Gαi1 and Gαi3 in NLGN3-induced signaling in MI.

## METHODS

### Data Availability

Detailed methods can be found in the Methods in Supplemental Material. Please see the Major Resources Table in the Supplemental Material.

### Generation of Gαi1/3 DKO mice

All animal experimentation was approved by the Institutional Review Board (IRB) of Soochow University (Suzhou, China). Generation of Gαi-KO mice by the CRISPR-Cas9 method was performed by GenePharma. Superovulation of C57/B6 donor female mice was performed by administering chorionic gonadotropin (CG) to pregnant mares, followed by the injection of 5 IU of human CG 48 h later. Female mice were mated with male mice and fertilized eggs were microinjected the next day. Microinjection of sgRNA (targeting mouse Gnai1 or Gnai3) and Cas9 mRNA was performed as described above[28]. Embryo-modified mice were born 19-20 days after microinjection. Newborn mice (F0) were genotyped at P7 and further characterized as Gnai1 or Gnai3 SKOs. Female chimeric Gnai+/-week 4-5 SKO mice were again mated with WT males and the newborn (P7) mice (G1) were genotyped. Several positive Gnai-SKO mice were identified, indicating the CRISPR-Cas9 genomic modifications had been integrated into germ cells and thus establishing the Gnai-SKO mouse line. The Gnai1/Gnai3-DKO mice were obtained by crossing Gnai1-SKO and Gnai3-SKO mice. Two mouse Gnai1 sgRNA sequences were tested: TCGACTTCGGAGACTCTGCT (target 1) and CCATCATTAGAGCCATGGGG (target 2). Two mouse Gnai3 sgRNA sequences were also tested: TTTTTAGGCGCTGGAGAATCTGG (target 1) and CATTGCAATCATACGAGCCATGG (target 2). In both cases, target 1 successfully induced Gnai SKO. Primers for genotyping were Gnai1: sense, GGTGAGTGAAGAGCCTACGG/ antisense, CACAGCGACTGGACCTCAAA; and Gnai3: sense, GGAGGGTTGCTTATGGAAT/ anti-sense, ACCTAACACTTCAAAAACAGA.

### Primary culture of ECs and Cardiac Fibroblasts

Primary neonatal (P1) mouse cardiac endothelial cells (MCECs) were isolated as described above[29]. Briefly, whole hearts were cut up, minced into small pieces, and digested with protease/collagenase II. The cells were then washed and incubated with magnetic beads conjugated with anti-CD31 antibody. Isolated cells were cultured in M199 medium with heparin, d-valine, IFCS, EGS, penicillin/streptomycin, and 10% fetal bovine serum. Purified ECs were stained with the EC marker CD31. Primary cells were used after 5 days of culture. Cardiac fibroblasts (CFs) were harvested from 1∼3-day-old neonatal mice. Hearts were rapidly excised and immersed in DMEM medium (10-013-cv, Corning, USA) containing 2% double antibody (pre-cooled), sheared to < 1 mm^3^, and digested in 0.25% trypsin (Thermo Fisher, USA) and 0.1% collagenase II (BioFroxx, Germany) at 37°C. After digestion, the cells were collected, centrifuged at 1000 rpm for 3 minutes, and then cultured in DMEM supplemented with 10% fetal bovine serum (Thermo Fisher, USA), 100 U/ml penicillin, and 100 μg/ml streptomycin (Thermo Fisher, USA) for 2 h. The medium was then changed to remove weakly adherent cells. Isolated and purified fibroblasts were further incubated in DMEM at 37°C in humidified air with 5% CO_2_ and 95% O_2_. Routine passages after 2 to 3 days P1-2 were used for experiments.

## RESULTS

### 1. Dynamic expression of NLGN3 after infarction

The serum levels of NLGN3 were determined by ELISA on the 7^th^ day after AMI in 20 acute MI patients and in 19 normal participants as controls. NLGN3 levels were significantly higher in the acute MI patients compared to the controls (Fig. 1A). Next, NLGN3 expression was analyzed in mouse hearts following permanent ligation of the left coronary artery. In this model, NLGN3 expression was found to be significantly upregulated in the early post-MI period (Fig. 1B, C).

**Figure 1.**
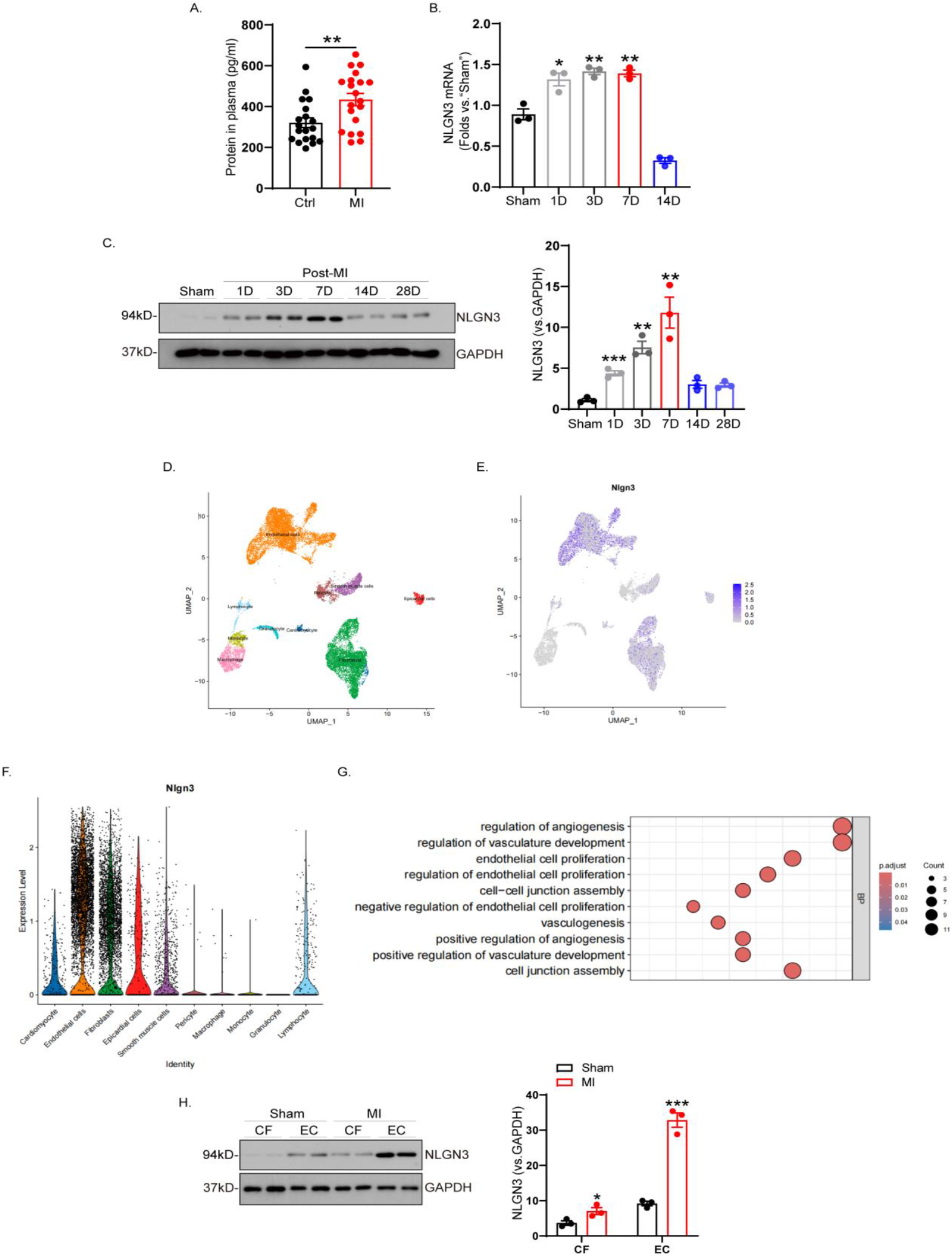
Myocardial infarction triggers the upregulation of NLGN3. A. NLGN3 levels in patient serum 7 days post-infarction (Ctrl, n=19; MI, n=20). B and C, mRNA (B) and protein levels (C) and quantitative analysis of NLGN3 in mouse hearts at 1, 3, 7, 14, and 28 days after myocardial infarction (MI) (n = 3 per group). D. Publicly available cardiac scRNA-seq data (GEO: #GSE153480) from MI and normal mice (sham) were visualized using uniform manifold approximation and projection (UMAP). E. Density plots show the expression and spatial distribution of NLGN3 in the heart, with the magnitude of the expression density on the right-hand side of the graph. F. Violin plot showing the scaled expression of NLGN3 in mouse heart snRNA-seq data across cell types as shown in Fig. E. G. Enrichment analysis showed that 48 co-expressed genes (CEGs) were associated with possible biological processes (BPs) of NLGN3 in the cardiac tissues of the MI mouse model. H. Western blotting of cardiac fibroblasts and ECs from sham-operated or MI mice, respectively. NLGN3 was predominantly expressed in cardiac ECs in isolated hearts. Data are expressed as the mean ± standard error of mean (SEM). (n = 3 per group). \**P*<0.05, \*\**P*<0.01, \*\*\**P*<0.001 vs “sham”

To identify the cell types with elevated NLGN3 expression, we analyzed publicly available single-cell RNA (scRNA) sequencing data (GEO: #GSE153480). Differential gene expression analysis between sham-operated and infarcted mice revealed that a total of 400 genes were significantly upregulated after infarction. This allowed exploration of the cellular localization of NLGN3 within the heart. Through dimensionality reduction, clustering, and annotation techniques, the individual cells throughout the heart were categorized into 10 distinct clusters (Fig. 1D). Subsequently, these clusters were visualized using uniform manifold approximation and projection (UMAP; Fig. 1E, F). As shown in Fig. 1F, NLGN3 was commonly expressed in all cell types within the mouse heart, with the highest expression in CFs and ECs.

Subsequently, primary CFs and ECs were extracted from infarcted mouse hearts and Western blotting was performed. This revealed that NLGN3 protein expression was significantly upregulated in ECs following infarction (Fig. 1G).

### 2. Silencing of NLGN3 in ECs Impedes Cardiac Function and Repair After Myocardial Infarction and Reduces Angiogenesis

To further examine the cardioprotective role of NLGN3 *in vivo*, MI surgery was performed on NLGN3 knockdown (AAV9-TIE1-shNLGN3) mice and on empty AAV9 control virus (AAV9-TIE1-sh-scr) mice. Cardiac function, cardiac remodeling, and angiogenesis were compared between the two groups (Fig. 2A). Three days after MI, cardiac function was significantly impaired in the NLGN3 knockdown group (Fig. 2B; Table S2). Moreover, the ischemic area was larger in NLGN3-cKD mice one day after MI (Fig. 2C).

**Figure 2.**
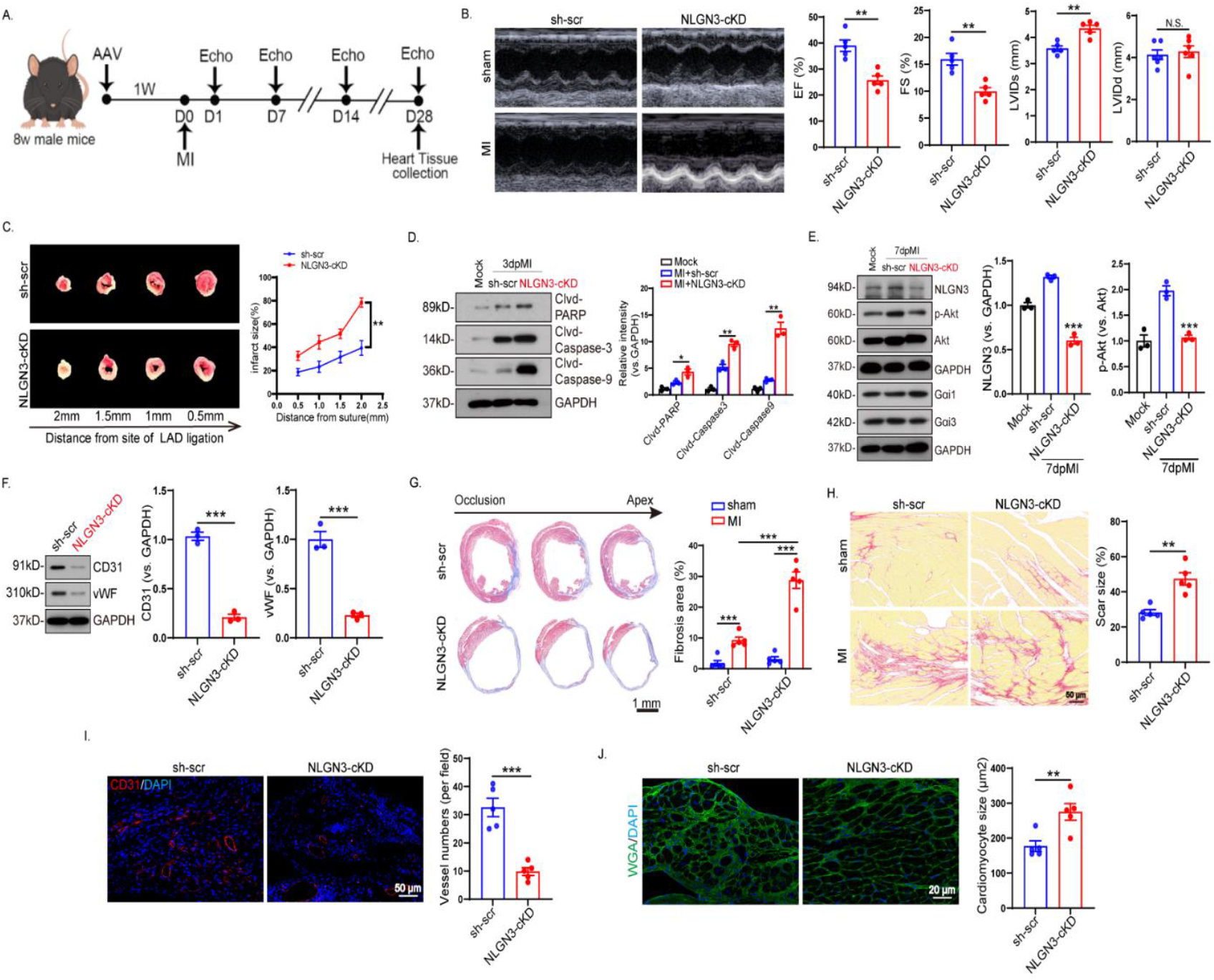
Cardiac EC silencing by NLGN3 exacerbates MI-induced ischemic heart injury in mice. A. Experimental design. NLGN3 shRNA-expressing AAV (NLGN3 shRNA-Tie1-AAV9, “NLGN3-cKD”) and the scrambled control shRNA-expressing adenovirus (“sh-scr”) were injected into mice, which were the subjected to the MI procedure 7 days later. B. The ischemic area was observed 24 h post-operatively by TTC staining. C. Echocardiography was performed to evaluate cardiac function at 72 h postoperatively. D, E and F. The infarcted area of the heart was isolated after the indicated time period and the specific protein expression was evaluated. G. Representative images and quantification of Masson trichrome staining to evaluate the infarct area in cKD and control mice (sh-scr) at 4 weeks post-MI. H. Representative images of the left ventricle of mice treated with scramble (sh-scr) and cKD were stained with Sirius red to show the degree of fibrosis. I. Representative CD31+ immunofluorescence images and quantification of infarcted cellular areas at 7 days post-MI. J. WGA staining to assess cardiomyocyte size at 4 weeks post-MI after virus injection. Data are expressed as the mean ± standard error of mean (SEM). (n = 5 per group). \**P*<0.05, \*\**P*<0.01, \*\*\**P*<0.001 vs “sh-scr”

Subsequently, Western blot assay was performed on isolated ischemic heart tissues from mice three and seven days after MI. Cardiac EC silencing of NLGN3 was found to enhance MI-induced apoptosis. Elevated levels of cleaved caspase-3, cleaved caspase-9, and cleaved PARP were observed in ischemic heart tissues from NLGN3-cKD mice (Fig. 2D). In addition, MI-induced cleavage and expression of NLGN3, and activation of Akt, were largely suppressed in the NLGN3-cKD group of mice (Fig. 2E). NLGN3 shRNA had no effect on Gαi1 and Gαi3 protein expression (Fig. 2E). We also extracted mouse heart tissue and used Western blot to evaluate the expression of CD31 and vWF after virus injection. The expression of both CD31 and vWF was lower compared to the sh-scr group (Fig. 2F).

Compared with AAV9-TIE1-sh-scr mice, NLGN3-cKD mice showed significantly more adverse remodeling 4 weeks after MI, with increased infarct size (Fig. 2G) and exacerbated cardiac fibrosis (Fig. 2H). Furthermore, EC-conditional knockdown of NLGN3 led to a significant reduction in the number of interstitial blood vessels in the peri-infarct area (Fig. 2I), and a larger cardiomyocyte area (Fig. 2J). Taken together, these results suggest that EC-specific knockdown of NLGN3 impairs cardiac function and exacerbates adverse pathological remodeling due to disrupted angiogenesis after MI.

### 3. Inhibition of NLGN3 cleavage by ADAM10 exacerbates ischemic heart injury in MI mice

NLGN3 is cleaved in neurons primarily by ADAM10. The ADAM10 inhibitor GI254023X (ADAM10i) was injected into the tail vein of mice 24 h before surgery for MI. After surgery, the mice were subjected to daily tail vein injections of ADAM10i (50 mg/kg body weight) or vehicle (0.9% NaCl solution) for 7 days (Fig. 3A).

**Figure 3.**
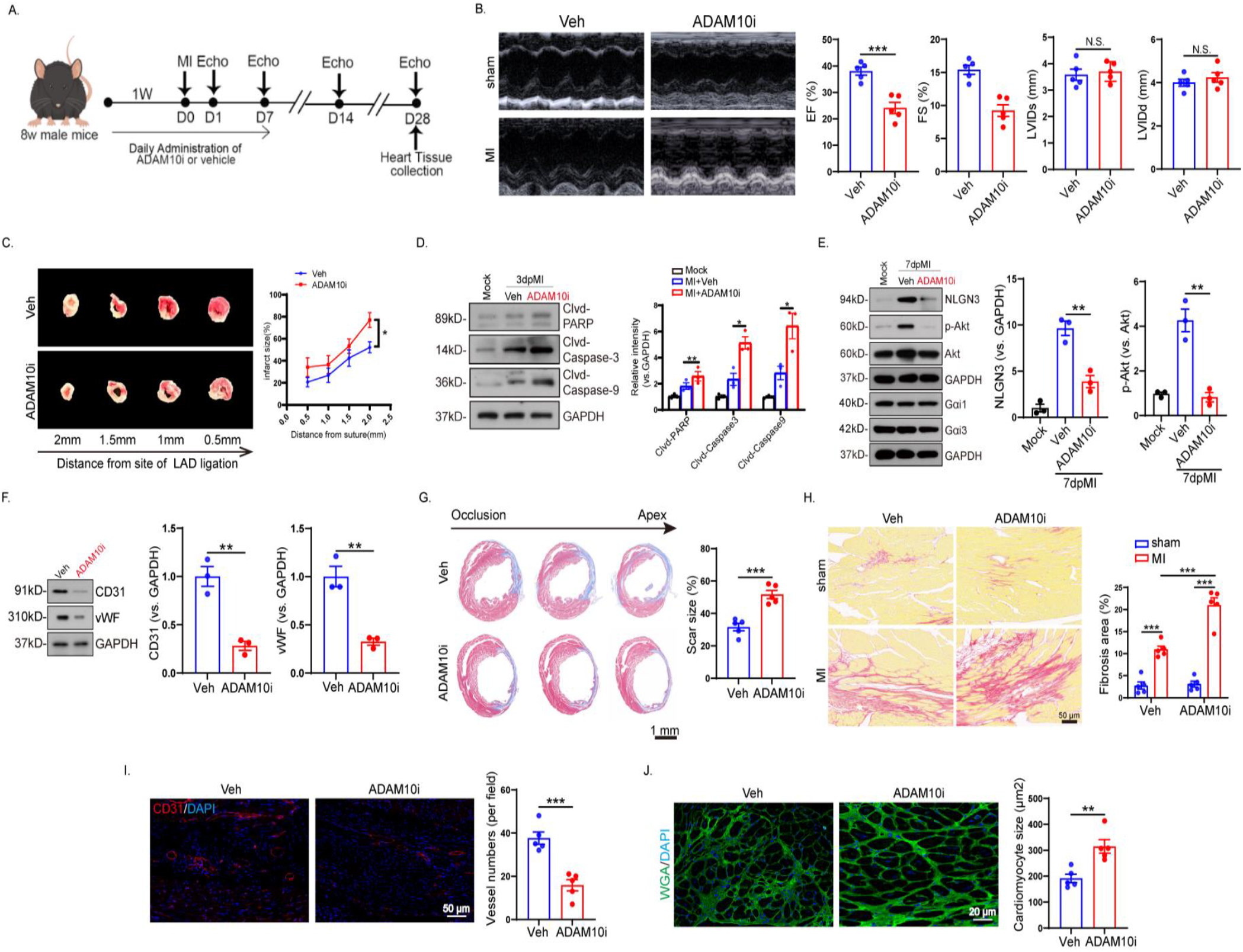
Inhibitor of ADAM10 inhibits NLGN3 cleavage and aggravates MI-induced ischemic cardiac injury in mice. A. Experimental design. ADAM10i (100 mg/kg) was injected daily into the tail vein for one week before and one week after MI surgery. B. Echocardiographic evaluation of cardiac function at 7 days post-MI in mice treated with vehicle or ADAM10i. C. The ischemic area was observed 24 h post-operatively using TTC staining. D, E and F. The infarcted area of the heart was isolated and the expression of the listed proteins was evaluated after the indicated time period. G. Representative images and quantification of Masson staining in vehicle-and ADAM10i-treated MI mice. H. Sirius red staining showing different degrees of fibrosis in the hearts of vehicle- and ADAM10i-treated mice, with quantification of the fibrosis. I. Detection of angiogenesis in the infarcted border area by staining for CD31 (green) and DAPI (blue). J. Representative WGA-stained images of myocardium in the peri-infarct zone of vehicle- and ADAM10i-treated mice hearts. Data are expressed as the mean ± standard error of the mean (SEM) (n = 5 per group). \**P*<0.05, \*\**P*<0.01, \*\*\**P*<0.001 vs “veh”

After 7 days of ADAM10i treatment, cardiac function was significantly decreased in the ADAM10i-treated mice compared with the vehicle group (Fig. 3B; Table S3). TTC staining showed an increase in the MI area one day after MI surgery (Fig. 3C). Pharmacological inhibition of NLGN3 by ADAM10i enhanced MI-induced apoptosis, and ischemic cardiac tissues showed significant increases in cleaved caspase-3, cleaved caspase-9, and cleaved PARP (Fig. 3D). Mice treated with ADAM10i showed a marked reduction in MI-induced cleavage and expression of NLGN3, and in the activation of Akt (Fig 3E). Compared to the vehicle control group, the expression levels of CD31 and vWF in the heart after ADAM10i injection were both reduced, as determined by Western blot analysis (Fig. 3F).

Four weeks after MI, the hearts of ADAM10i mice showed increased scar size (Fig. 3G), increased fibrosis (Fig. 3H), and a decreased number of vessels in the peri-infarct zone (Fig. 3I) compared with the control group. Furthermore, pharmacologic inhibition of NLGN3 by ADAM10i increased the size of cardiomyocytes (Fig. 3J), thereby worsening pathological remodeling. These results suggest that ADAM10 inhibitors impaired the MI-induced cleavage and secretion of NLGN3, and aggravated ischemic cardiac injury in mice.

### 4. NLGN3 ameliorates hypoxia-induced apoptosis in cardiac ECs

NLGN3 expression in primary cardiac ECs showed the most increase after 24 h of hypoxic stimulation (Fig. 4A). Enhanced cleavage of Caspase-3, Caspase-9, and PARP was detected in hypoxia-stimulated cardiac ECs, but these apoptosis markers were reduced after NLGN3 pretreatment (Fig. 4B). Hypoxic stimulation induced marked apoptotic activation in cardiac ECs, a significant increase in the proportion of TUNEL-positive nuclei (Fig. 4C), and an increased rate of apoptosis as detected by Annexin V/PI flow cytometry (Fig. 4D). The pro-apoptotic effect of hypoxia was largely attenuated by pretreatment with NLGN3 (Fig. 4C, D). In summary, NLGN3 ameliorated hypoxia-induced apoptosis in cardiac ECs.

**Figure 4.**
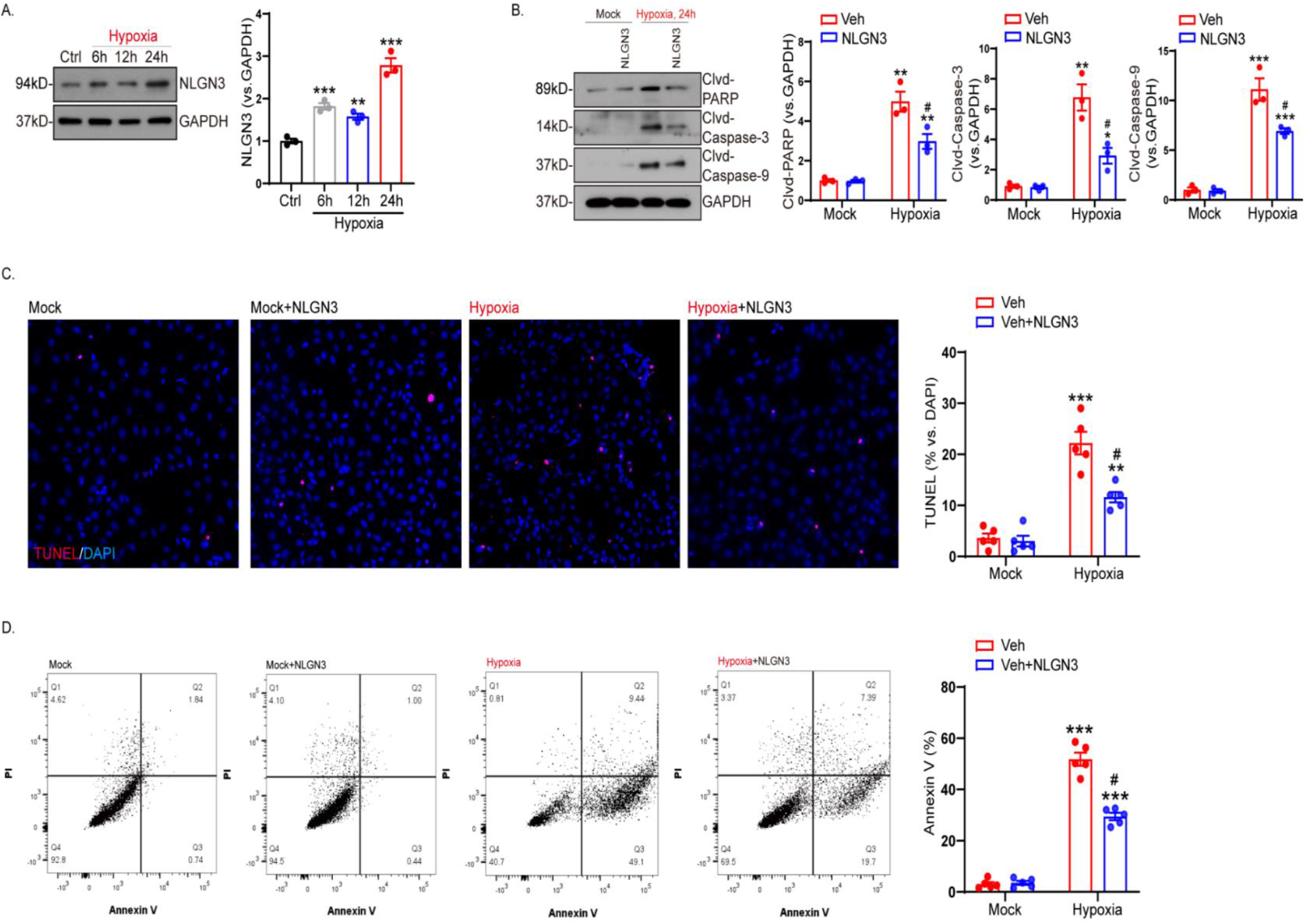
NLGN3 ameliorates hypoxia-induced apoptosis in cardiac ECs. A. Cardiac ECs were hypoxia-treated for 0, 6, 12, and 24 h, and NLGN3 protein expression was then detected by Western blotting. B. Cardiac metaplastic ECs were pretreated with NLGN3 (50 ng/mL) or control (PBS, “veh”) for 10 min after hypoxia treatment for 24 h. Expression of the listed proteins was evaluated by Western blotting. C and D. Apoptosis was detected by nuclear TUNEL staining (C) and Annexin V-PI FACS (D), and the results quantified. “Mock” indicates mock processing. Western blot data are representative of three replicate experiments. Data are expressed as the mean ± standard error of the mean (SEM). (n = 3 or 5 per group). \**P*<0.05, \*\**P*<0.01, \*\*\**P*<0.001 vs “mock”. #p<0.05 vs “veh”

### 5. Gαi1 and Gαi3 mediate NLGN3-induced activation of Akt signaling in ECs

Cardiac ECs were treated with NLGN3 at progressively increasing concentrations from 25 to 200 ng/ml. NLGN3 was found to increase the phosphorylation levels of Akt (Ser-473), S6, and mTOR in a concentration-dependent manner (Fig. 5A). NLGN3-induced activation of Akt-mTOR was highest at 50 ng/mL and after 10 min of treatment (Fig. 5B). These conditions were therefore used in the following experiments.

**Figure 5.**
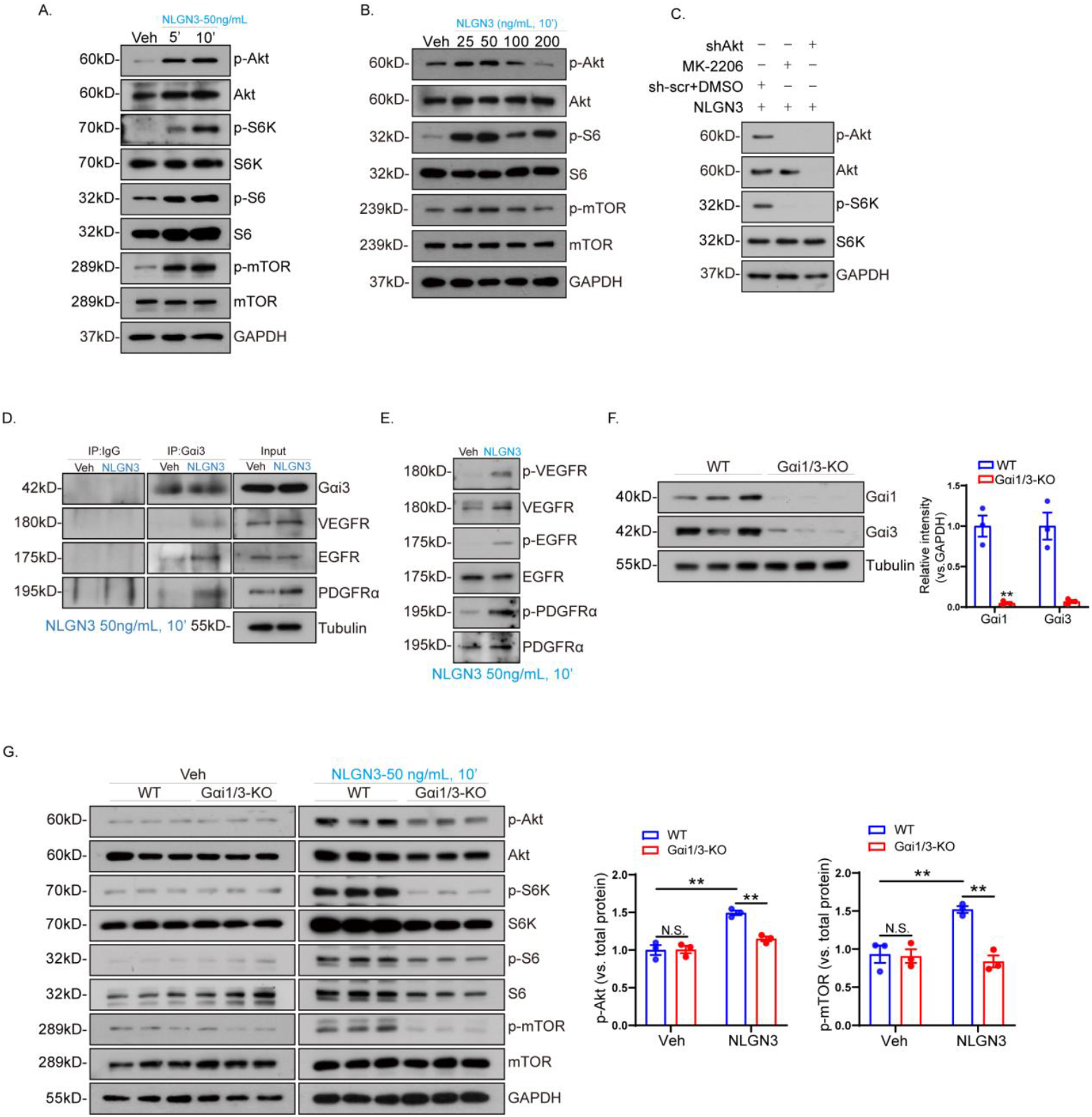
Gαi1 and Gαi3 mediate NLGN3-induced Akt signaling activation in cardiac ECs. A and B. Cardiac primary ECs were treated with NLGN3 or control (PBS, “veh”) for 10 min and the expression of the listed proteins was then evaluated by Western blotting. C. ECs were pretreated with MK-2206 (10 μM, 30 min) or stably transduced with Akt shRNA (“shAkt”), followed by treatment with NLGN3 (50 ng/mL, 10 min). Expression of the listed proteins was then evaluated. D and E. Cardiac ECs were treated with NLGN3 (50 ng/mL) for 10 min. Expression of the listed proteins was then detected by Western blotting. Interactions between RTKs (PDGFRα, EGFR, VEGFR) and Gαi3 were detected by immunoprecipitation assay (D), and phosphorylated protein expression of PDGFRα, EGFR, and VEGFR by Western blotting (E). F and G. WT and Gαi1/3-KO primary mouse cardiac ECs were first treated with NLGN3 and expression of the listed proteins was then evaluated by Western blotting. Data are expressed as the mean ± standard error of the mean (SEM). (n = 5 per group). \*\**P*<0.01 vs “WT” and “Veh”. Each experiment was repeated 3 times, with similar results obtained.

The Akt-specific inhibitor MK-2206 and Akt shRNA lentiviral particles were used to block Akt activation. NLGN3-induced phosphorylation of S6K was almost completely blocked by MK-2206 and shAkt in cardiac ECs (Fig. 5C).

Co-immunoprecipitation experiments showed that Gαi1/3 was associated with NLGN3-activated RTKs in cardiac ECs (Fig. 5D). Furthermore, NLGN3 was found to induce the phosphorylation of multiple RTKs (e.g., VEGFR, EGFR) in cardiac ECs (Fig. 5E). Additionally, WT and Gαi1/3-KO primary mouse cardiac ECs were extracted and pretreated with 50 ng/mL NLGN3 for 10 minutes. Western blotting showed that NLGN3 significantly enhanced downstream signaling in ECs from WT mice, whereas no significant changes in downstream signaling were observed in ECs from knockout mice (Fig. 5F, G).

### 6. Gαi1/3 plays a key role in NLGN3-induced EC proliferation and tube formation

The expression of Gαi1 and Gαi3 proteins was significantly reduced in stable HCAECs with the aforementioned Gαi1 and Gαi3 shRNAs compared to scrambled shRNAs (“sh-scr”), while phosphorylation of the downstream signals Akt, S6K, S6 and mTOR was barely detectable (Fig. 6A). However, the phosphorylation levels of these downstream signals were significantly up-regulated after NLGN3 treatment, but not in HCAECs with knockdown of Gαi1 and Gαi3 (Fig. 6B).

**Figure 6.**
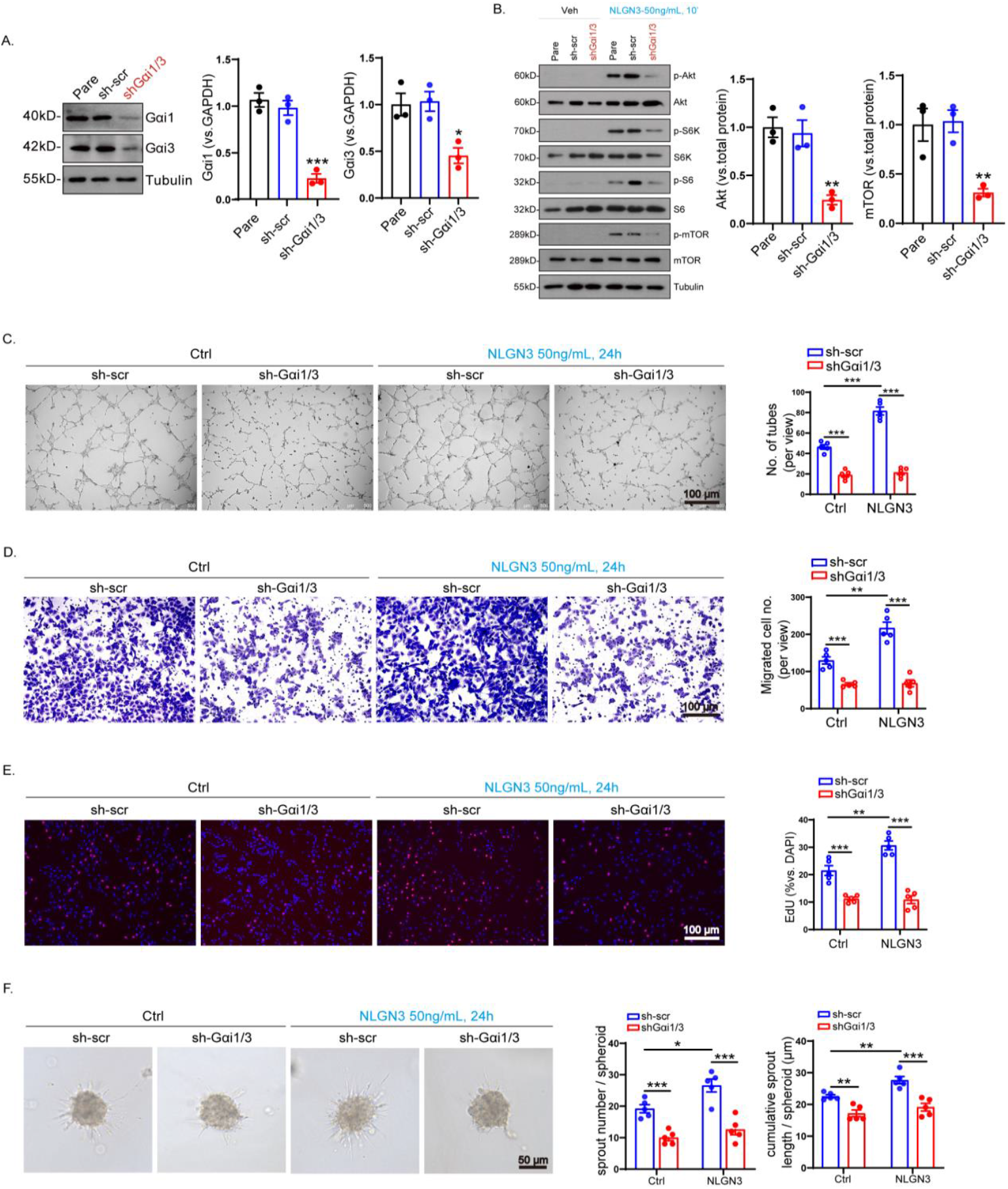
Gαi1/3 silencing blocks NLGN3-induced HCAEC angiogenesis *in vitro*. A. The graphs show Gαi1/3 protein expression in human coronary artery endothelial cells (HCAEC) stabilized by Gαi1/3 shRNA, scramble shRNA (“sh-scr”), or parental control (“pare”). B and H. Three kinds of cells were treated with NLGN3 (50 ng/mL for 10 min) and expression of the listed proteins was evaluated by Western blotting. C, D, E, F. The same number of HCAECs were placed in complete culture medium and cultured for the indicated times. Cell proliferation was evaluated *in vitro* by measuring the binding of EdU to the nucleus (C), migration with the “Transwell” assay (D), capillary tube formation (E), and sprout formation (F). Data are expressed as mean ± standard error of the mean (SEM). (n = 3 or 5 per group). \**P*<0.05, \*\**P*<0.01, \*\*\**P*<0.001 vs “sh-scr” or “ctrl”. Western blot experiments were repeated 3 times and similar results were obtained.

The proliferation, migration, tube formation and outgrowth assays were performed on HCAECs after 24 h of pretreatment with NLGN3. Compared to the sh-scr group, NLGN3 accelerated *in vitro* tube formation (Fig. 6C) and migration (Fig. 6D) of HCAECs with Gαi1 and Gαi3 shRNAs. The percentage of EDU-positive nuclei was also increased (Fig. 6E), as well as the number and length of sprouts (Fig. 6F). However, tube formation (Fig. 6C), cell migration (Fig. 6D), proliferation (Fig. 6E), and sprout capacity (Fig. 6F) were reduced in HCAECs with knockdown of Gαi1/3, which was not significantly altered by pretreatment with NLGN3. The above results indicate that NLGN3 promotes angiogenesis by HCAECs *in vitro*, but has no significant effect after knockdown of Gαi1/3, suggesting that Gαi1/3 is a key channel protein downstream of NLGN3.

### 7. Gαi1 and Gαi3 deficiency negatively affects cardiac function, remodeling and angiogenesis after myocardial infarction

We established a MI model in wild-type (WT) and Gαi1/3 knockout (Gαi1/3-KO) mice and tested cardiac function 4 weeks after MI (Fig. 7A). Gαi1/3 deficiency significantly impaired cardiac function. Post-MI, the EF and FS gradually decreased, and LVIDd and LVIDs gradually increased. Furthermore, these changes were more pronounced in the Gαi1/3-KO group (Fig. 7B; Table S3). In addition, Masson staining showed the infarct area was larger in Gαi1/3-KO mice than in the control group (Fig. 7C). Sirius red staining was also performed on the heart tissue of mice to assess the degree of cardiac fibrosis. Following MI, cardiac fibrosis was more extensive in the infarct zone of Gαi1/3-KO mice compared with WT mice (Fig. 7D).

**Figure 7.**
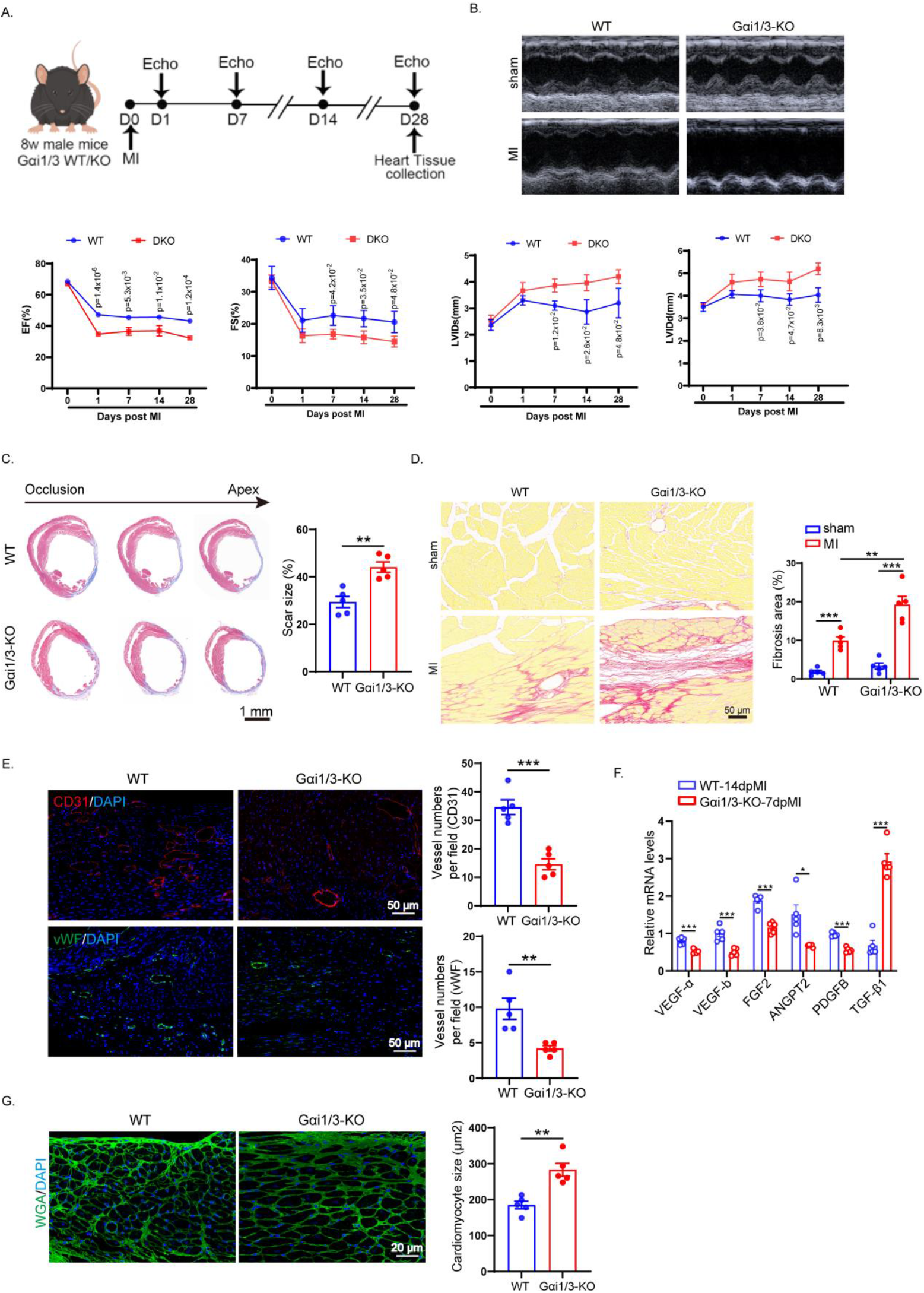
Gαi1/3 knockout (KO) worsens cardiac function, remodeling, and angiogenesis after myocardial infarction (MI) in mice. A. Schedule of animal experiments. Adult (8-week) WT or Gαi1/3-KO mice were subjected to MI or sham operation. Hearts were removed 4 weeks after myocardial infarction (4wpMI). B. Echocardiographic evaluation of cardiac function, with representative images of LVEF, LVFS, LVIDD, and LVIDS analyses. C. Representative images and quantification of Masson’s trichrome staining to assess infarct size in mice 4 at wpMI. D. Representative images and quantitative analysis of Sirius red staining for myocardial fibrosis in the infarct border zone of 4wpMI hearts. E. Representative immunofluorescence images and statistical analysis of cardiac vessel counts. The EC markers vWF and CD31 were used to stain blood vessels at 4wpMI and 7 dpMI, respectively, and DAPI to stain nuclei. F. Heart tissues were collected from WT and Gαi1/3-KO mice 7 days after MI. Relative mRNA levels of angiogenic factors were quantified by qRT-PCR. G. Representative WGA staining images and quantification of cardiomyocytes in the border zone at 4 wpMI. Data are expressed as the mean ± standard error of the mean (SEM). (n = 5 per group). \**P*<0.05, \*\**P*<0.01, \*\*\**P*<0.001 vs “WT”

Subsequently, vWF/CD31 staining was performed to assess angiogenesis. Compared with the WT group, vascular numbers were significantly reduced in Gαi1/3-KO hearts at 7 days and 4 weeks post-MI (Fig. 7E). The levels of angiogenesis-related factors (VEGF-a, VEGF-beta, FGF2, ANGPT2, PDGFB, TGF-beta1) were all increased 7 days post-infarction, and to a greater extent in the WT group (Fig. 7F).

To further assess cardiac remodeling, WGA staining was performed to observe changes in the size of cardiomyocytes after MI. The results showed that cardiomyocytes from both groups of mice increased in size at 4 weeks post-MI, but more so in Gαi1/3-KO mice (Fig. 7G). Taken together, these results suggest that Gαi1/3 plays an important role in cardiac injury and regeneration in adult mice.

### 8. Gαi1 and Gαi3 Deficiency are Involved in Early Cardiac Injury in Mice with Myocardial Infarction

Cardiac function, fibrosis, and remodeling are all long-term changes in the heart after MI. We next investigated the role of Gαi1/3 in the early stages of MI. At one day post-MI, the ischemic area was assessed by 2,3,5-triphenyltetrazolium chloride (TTC) staining. This revealed a significant increase in the size of the ischemic area in Gαi1/3-KO mice (Fig. S1A). Consistent with the increase in acute ischemic injury, serum LDH levels were significantly higher 3 days after surgery in Gαi1/3-KO mice compared with the WT group (Fig. S1B). Additionally, TUNEL staining showed a greater increase in apoptosis in the hearts of Gαi1/3-KO mice 3 days after MI (Fig. S1C). CD68 staining was used to observe macrophage infiltration in cardiac tissues from both groups of mice. This revealed a slight increase in infiltration in Gαi1/3-KO mice compared with the WT group (Fig. S1D). Primary cardiac ECs were isolated from WT and Gαi1/3-KO mice and treated by hypoxia for 4 h. A significant increase in apoptosis was detected in the Gαi1/3-KO mice by Annexin V/PI flow cytometry (Fig. S1E). In conclusion, these results suggest that Gαi1/3 deficiency may affect EC function and aggravate cardiac injury following MI.

## 4. DISCUSSION

This study found that NLGN3 plays a critical role in maintaining endothelial integrity and improving cardiac function after MI. Decreased expression of NLGN3 in ECs leads to structural and functional cardiac defects, increased apoptosis in the early infarct phase, and reduced vascular density in the infarct border zone. NLGN3-induced Gαi1/3 promotes EC proliferation, migration, and tube formation through the PI3K-Akt-mTOR pathway, suggesting that Gαi1/3 is a key channel protein that mediates NLGN3 downstream signaling (Fig. 8).

**Figure 8.**
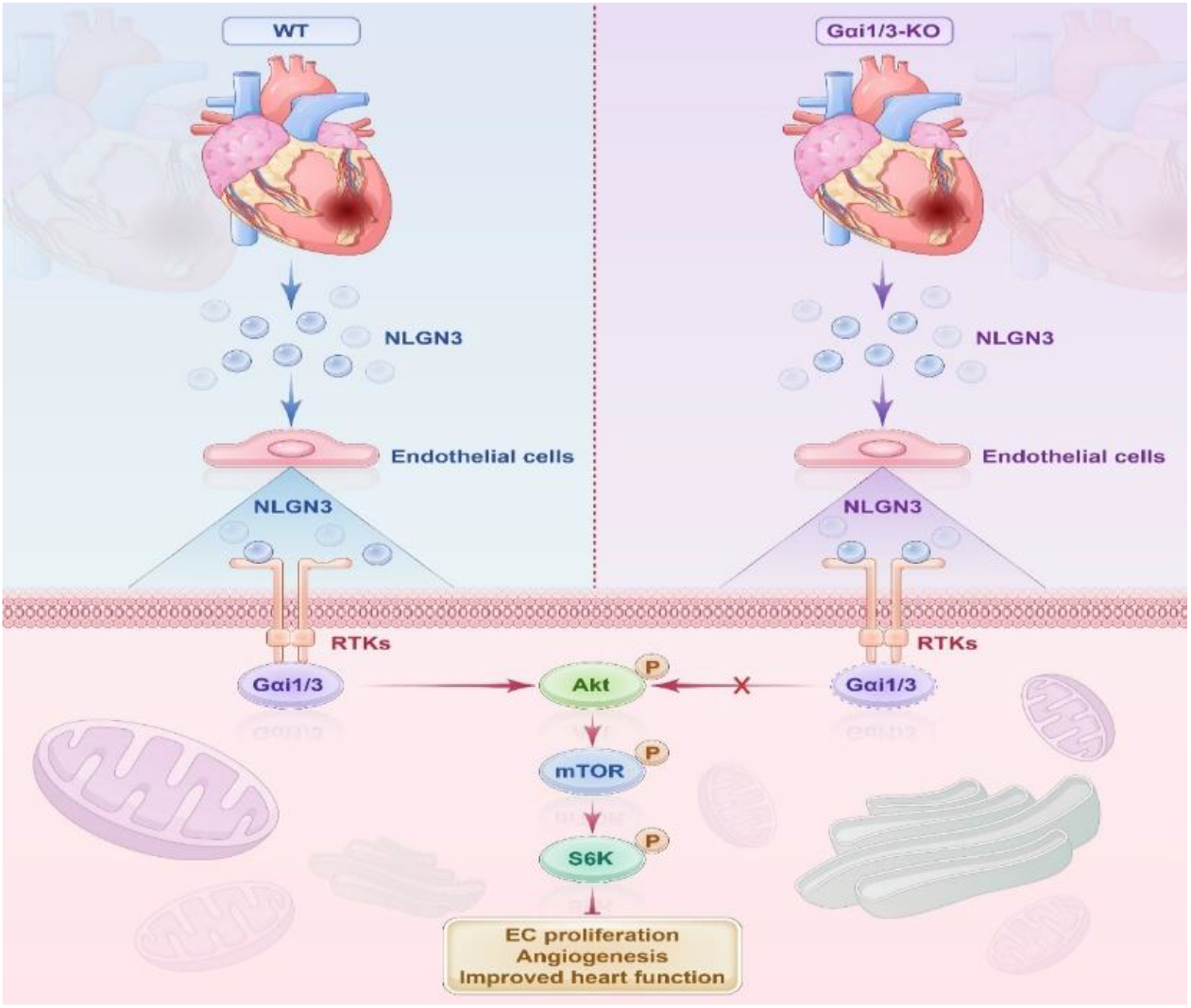
Schematic diagram showing how NLGN3 provides protection against myocardial infarction (MI).

As a member of the neural synaptic protein family, NLGN3 displays a complex pattern of selectivity, with effects that are specific to excitatory or inhibitory synapses, and which acts in a context-dependent manner[30-33]. NLGN1 has been shown to regulate vascular morphogenesis *in vitro* and in the mouse retina[34], while NLGN2 is expressed in vascular ECs and may regulate angiogenesis by modulating the release of vascular regulatory factors[35]. However, to our knowledge the effects of NLGN3 on the vasculature have not been reported previously. Our clinical study found that the plasma concentration of NLGN3 is significantly increased in patients with coronary artery occlusion. In addition, NLGN3 expression increases in the mouse heart after MI. Further analysis revealed that NLGN3 is highly expressed in ECs of the heart. Together, the above results indicate that NLGN3 is involved in the pathological changes that occur after MI, and that its mechanism of action may involve regulating the function of myocardial blood vessels.

Angiogenesis is an important mechanism for improving MI. The promotion of angiogenesis, as measured by myocardial capillary density, can improve heart failure after MI and improve myocardial remodeling. This study found that an exogenous increase in NLGN3 significantly enhanced the tube-forming ability of both human coronary ECs and primary mouse ECs. Moreover, pretreatment with NLGN3 also significantly reduced the rate of apoptosis after hypoxia induction. Furthermore, EC-specific knockdown of NLGN3 following mouse heart surgery resulted in decreased vascular density during both early and late infarction, suggesting an important role for NLGN3 in post-infarction angiogenesis. Experiments with a specific NLGN3 blocker also showed that NLGN3 plays a key role in angiogenesis following MI. ADAM10 is one of the most important proteases with specific cell surface protein regulatory activity[36]. In mice subjected to experimental infarction, administration of the ADAM10 inhibitor GI254023X via a minipump for as little as 3 days showed therapeutic potential by improving survival and enhancing cardiac function[37]. Importantly, inhibition of ADAM10 was shown to cause a dose-dependent increase in full-length NLGN3 in brain slice lysates, along with decreased cleavage of NLGN3 and reduced NLGN3 shedding[17]. We therefore injected GI254023X into the tail vein of mice to reduce NLGN3 secretion. Mice injected with ADAM10i had more severe post-infarction cardiac injury and poorer cardiac function, consistent with the *in vivo* knockdown of NLGN3.

The PI3K-Akt-mTOR pathway has been widely implicated in cardiac pathophysiology. Activation of this pathway in a rat diabetic myocardial I/R model can alter the dynamic balance between mitochondrial fusion and fission, as well as improving cardiac hemodynamic parameters[38]. Cardioprotective effects following activation of the PI3K/AKT/mTOR pathway were also observed in donor hearts exposed to prolonged cold ischemia[39]. More recently, activation of this pathway was shown to promote neovascularization and contribute to the growth of tumor tissue[40]. Neovascularization and the promotion of angiogenesis in myocardial tissue after MI is considered to be an effective treatment strategy that restores blood flow to the ischemic and hypoxic tissue. The PI3K/Akt/mTOR signaling pathway plays a crucial role in the formation of normal blood vessels. The downstream effectors HIF-1α, eNOS, VEGF-A and forkhead O transcription factor (FOXO) are involved in the generation and maintenance of blood vessels after MI[41]. Activation of the PI3K/AKT pathway in HCAECs can trigger angiogenesis in MI hearts and increase the density of CD31-positive vessels[42]. Consistent with previous studies, we found that treatment with NLGN3 (50 ng/ml) for 10 min can increase the expression of p-akt in HCAECs.

Therefore, the findings of our study suggest that the mechanism by which NGLN3 protects against myocardial failure after MI may be through the promotion of myocardial angiogenesis following activation of the PI3K-Akt-mTOR pathway.

Gαi1/3 expression has an important role in NLGN3-induced Akt-mTORC1 activation[18, 43]. In the current study, co-immunoprecipitation experiments with cardiac ECs showed that Gαi1/3 was associated with NLGN3-activated RTKs. Numerous studies have shown that Gαi1 and Gαi3 play important roles in angiogenesis and are key channel proteins that mediate downstream signaling[23, 44-47]. However, more exploration is needed into whether the activation of Gαi1/3-Akt signaling by NLGN3 contributes to angiogenesis after MI. Our experiments revealed that the expression of both Gαi1 and Gαi3 were upregulated after MI. Because angiogenesis is an important part of the post-infarction cardiac repair process, we also investigated the roles of Gαi1 and Gαi3 in post-infarction angiogenesis. As expected, we found that angiogenic capacity was reduced in both infarcted mouse hearts with knockout of Gαi1 and Gαi3, and in human cardiac ECs with knockdown of Gαi1 and Gαi3. Our findings indicate that NLGN3 activates signaling downstream of Gαi1/3 proteins in the heart by binding to RTKs, thereby promoting EC proliferation and tube formation. These results highlight the important role of NLGN3 in angiogenesis and in the improvement of heart function after MI.

This study has several limitations. First, NLGN3 is a neurosecretory protein. Our study found that plasma concentrations of NLGN3 were increased in patients with MI, and that specific silencing of NLGN3 in coronary ECs exacerbates heart failure after MI. Although similar findings were also obtained by administering the NLGN3 inhibitor ADAM10, we did not directly observe the effect of increasing plasma NLGN3 concentration on MI. Second, our study found that NLGN3 decreased 14 days after MI. We did not further explore whether the mechanism for the initial increase in NLGN3 was due to early activation of neural components during MI. Third, our previous research showed that NLGN3 is a neuroendocrine protein expressed in myocardial fibroblasts and ECs, but we also observed myocardial cell hypertrophy during the pathological process of MI. Additional studies are required to determine whether NLGN3 is involved in the pathological changes found in myocardial cells. Finally, our research focused on the mechanism of angiogenesis in MI. We found that NLGN3 regulates angiogenesis by activating the PI3K-Akt-mTOR pathway, but further research is needed to determine whether other pathways are involved in this regulation.

In conclusion, we conducted a comprehensive functional investigation on the role of NLGN3 in the progress of MI. Based on our findings, we propose a plausible molecular and cellular mechanism involving angiogenesis. The highest expression of NLGN3 was observed in cardiac ECs in the mouse heart. We provide evidence that Gαi1 and Gαi3 are the most likely target genes for NLGN3 in the regulation of HCAEC angiogenesis. Based on the exploration of tube formation, migration, proliferation, apoptosis and sprout capacity by HCAECs, we propose a mechanism involving, at least partially, activation of the PI3K/AKT/mTOR pathway. This work presents a molecular biological mechanism to explain the protective effect of NLGN3 after MI.

## Acknowledgments

We express our gratitude to all staff members of the Department of Cardiology at the Second Affiliated Hospital of Soochow University for their contributions to data collection.

## Non-standard Abbreviations and Acronyms

NLGN3: neuroligin-3
ADAM10i: a disintegrin and metalloproteinase 10 inhibitor
Gαi1/3: the inhibitory α subunit 1/3 of G proteins
RTKs: receptor tyrosine kinases
EGFR: epidermal growth factor receptor
PDGFR: platelet-derived growth factor receptor
VEGFR: vascular endothelial growth factor receptor
HCAEC: human coronary artery endothelial cell
HUVEC: human umbilical vein endothelial cell
MCEC: mouse cardiac endothelial cell
cKD: conditional endothelial cell knockdown
MI: myocardial infarction
WT: wild type
WGA: wheat germ agglutinin

